# Sensitivity of human sweet taste receptor subunits T1R2 and T1R3 to activation by glucose enantiomers

**DOI:** 10.1101/2022.12.19.520966

**Authors:** Nitzan Dubovski, Yaron Ben-Shoshan Galezcki, Einav Malach, Masha Y. Niv

**Affiliations:** The Institute of Biochemistry, Food Science and Nutrition, Robert H. Smith Faculty of Agriculture, Food and Environment, The Hebrew University of Jerusalem, P.O. Box 12, Rehovot 76100, Israel

## Abstract

We have previously shown that L-glucose, the non-caloric enantiomer of D-glucose, activates the human sweet taste receptor T1R2/T1R3 transiently expressed in HEK293T cells. Here we show that D- and L-glucose can also activate T1R2 and T1R3 expressed without the counterpart monomer. Serine mutation to alanine in residue 147 in the binding site of T1R3 VFT domain, completely abolishes T1R3S147A activation by either L or D-glucose, while T1R2/T1R3S147A responds in the same way as T1R2 expressed without its counterpart. We further show that the original T1R2 reference sequence (NM_152232.1) is less sensitive by almost an order of magnitude than reference sequence at the time this study was performed (NM_152232.4). We find that out of the four differing positions, it is the R317G in the VFT domain of T1R2, that is responsible for this effect in-vitro. It is important both for practical assay sensitivity, and because glycine is found in this position in ∼20% of the world population. While the effects of the mutations and of the partial transfections were similar for D and L enantiomers, their dose response curves remained distinct, with L-glucose reaching an early plateau.

## Introduction

The primary taste receptor for sweet sensing has been identified as a heterodimer of two Class C GPCRs-T1R2 and T1R3 (Li X et al., 2002). Family C GPCRs are obligate dimers, forming either homo- or hetero-dimers (Kniazeff J et al., 2011). The canonical sweet taste receptor in the human body acts as a T1R2/T1R3 heterodimer (Nelson G et al., 2001), and expression of both subunits has been detected in the oral cavity (Archer NS et al., 2016; Xi R et al., 2022) as well as in the stomach and intestine (Jang HJ et al., 2007). However, there is evidence for the function of T1R3/T1R3 homodimers in different cells in the human gastro intestinal (GI) tract (Buchanan KL et al., 2022; Li X, 2009; Margolskee RF et al., 2007; Ohtsu Y et al., 2014; Zhao GQ et al., 2003). Behavioral studies on T1R3 or T1R2 knock-out mice display responses to sugars, which show that T1R2 and mainly T1R3 could potentially form functional homodimers in mice (Treesukosol Y et al., 2011).

Studies of the sweet taste receptor have mapped multiple binding sites in different parts of the two monomers of the receptor. For instance, the venus fly trap (VFT) domain of T1R2 is the binding site of artificial sweeteners such as acesulfame-potassium, aspartame, saccharin and neotame (Masuda K et al., 2012), while both T1R2 and T1R3 VFT domain bind neutral sugars such as sucrose and glucose, as well as sucralose (Nie Y et al., 2005). The cysteine rich domain (CRD) of T1R3 binds sweet proteins such as brazzein and thaumatin (Masuda K et al., 2012). The transmembrane domain (TMD) of T1R3 was identified as the binding site of cyclamate (Xu H et al., 2004) and neohesperidin dihydrochalcone (NHDC) (Winnig M et al., 2007), and the sweet taste inhibitor lactisole (Jiang P et al., 2005), while the TMD of T1R2 is the binding site of some other sweeteners, such as S-819 and P-4000 (DuBois GE, 2016).

T1R2 and T1R3 are considered as some of the most polymorphic genes in the human genome (Chamoun E et al., 2018), with most of the single nucleotide polymorphisms (SNPs) are located on the VFT domain and TMD, where the ligands bind (Smith NJ et al., 2021). There is some evidence for the impact of T1R2 and T1R3 SNPs on eating behavior and sweet consumption. For example, the genetic variations S9C and I191V in T1R2 were found to be associated with sweet consumption and eating behavior (Eny KM et al., 2010; Haznedaroglu E et al., 2015; Kulkarni GV et al., 2013; Pioltine MB et al., 2018). In another study that evaluated the effect of the I191V variant of T1R2 on glucose homeostasis in healthy individuals, the valine allele was shown to correlate with a partial loss of function of the monomer, and reduced glucose excursions in the oral glucose tolerance test (Serrano J et al., 2021). However, in all these studies, the participants were genotyped for the presence of the SNPs in residues 9 and 191 only, and a full T1R2 sequencing was not performed. Similarly, SNPs on non-coding regions of T1R3 were also found to be associated with sugar consumption (Fushan AA et al., 2009).

Chirality is a fundamental property of molecular asymmetry that has significance in biological recognition. A molecule with a single chiral center has two enantiomeric forms, which are mirror images of each other. L-glucose is the enantiomer of D-glucose. However, it is not metabolized and hence considered as a non-caloric sugar (Levin GV et al., 1995). In our previous study, we have proved for the first time that L-glucose sweetness is mediated by T1R2/T1R3 (Dubovski N, Y Ben Shoshan-Galeczki, et al., 2022). Here we studied the contribution of each of the sweet receptor subunits, T1R2 and T1R3, in D- and L-glucose recognition, and evaluated the effects of SNPs found in the current vs original reference sequences of T1R2. Finally, we have shown the critical importance of T1R3 residue 147 for activation by both L-glucose and D-glucose.

## Results

In our previous study (Dubovski N, Y Ben Shoshan-Galeczki, et al., 2022) we used a T1R2 plasmid received from colleagues several years ago. Following recommendations from Smith et al (Smith NJ et al., 2021), we sequenced the plasmid and found that this is the original reference sequence (RefSeq) (NM_152232.1), and contains 4 SNPs compared to the updated release RefSeq of T1R2 (NM_152232.4, which we will refer to as wild type, WT). Each of these SNPs have a frequency of at least 1% in the population. Three of the SNPs-I191V, R317G and I486V are located on the VFT domain of T1R2 (see Figure 1), while S9C is located in the N-terminal signal-peptide.

**Figure 1:**
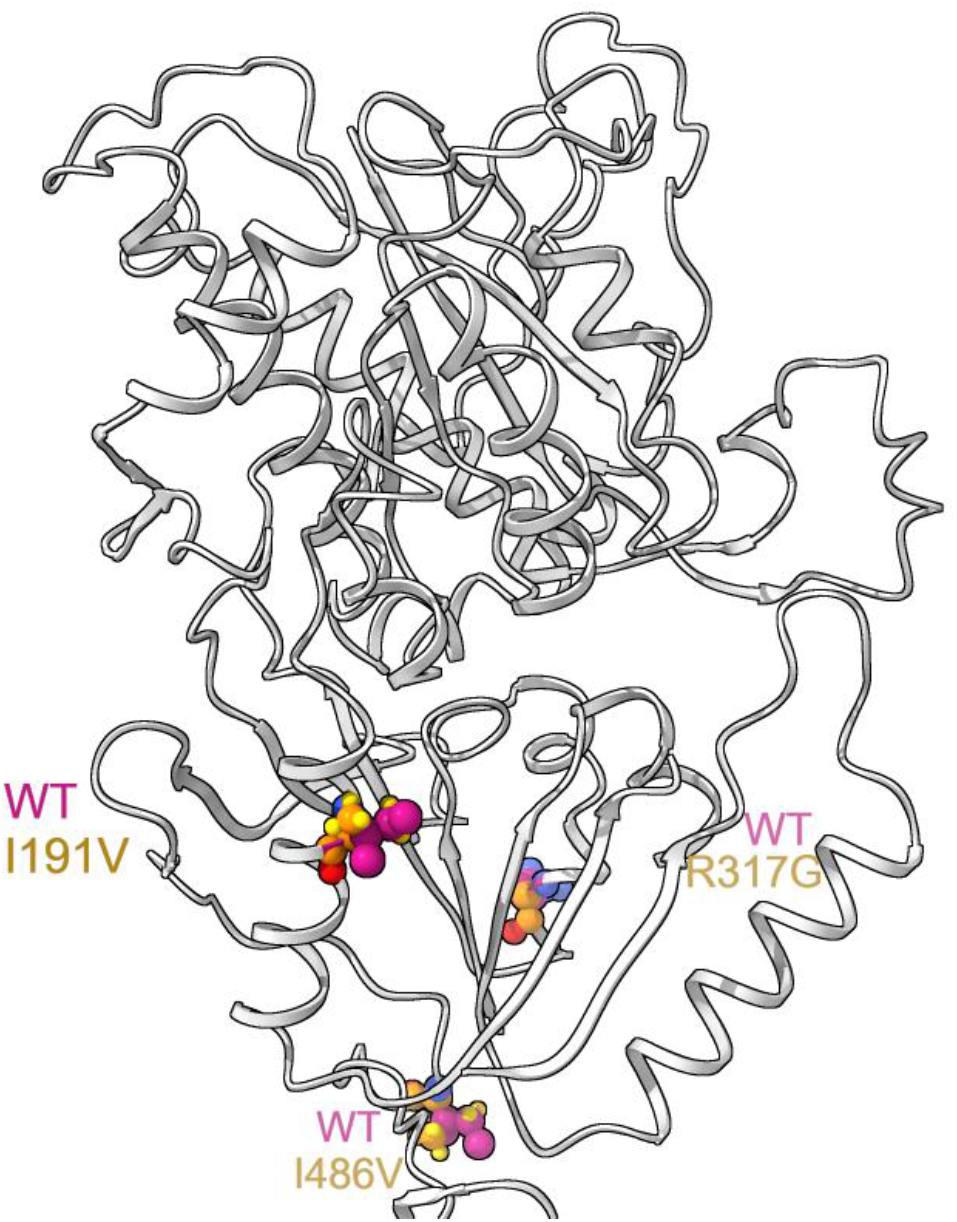
Visualization of T1R2 SNPs varying between the old (RefSeq NM_152232.1) and current (RefSeq NM_152232.4, WT) sequences, on the VFT domain.

In order to understand whether these SNPs affect the sensitivity of activation by L-glucose, we compared the dose response curves of HEK293T cells, transfected with either the “new WT” T1R2 (mentioned as T1R2) or with the original T1R2S9C,I191V,R317G,I486V, along with T1R3. Indeed, as shown in Figure 2A, we observed differences in both potency and efficacy: in transfection with WT T1R2 sequence, higher E_max_ (100%) and lower EC_50_ (0.002 M) were obtained, compared to the transfection with T1R2S9C,I191V,R317G,I486V (E_max_= 50%, EC_50_= 0.015 M).

**Figure 2:**
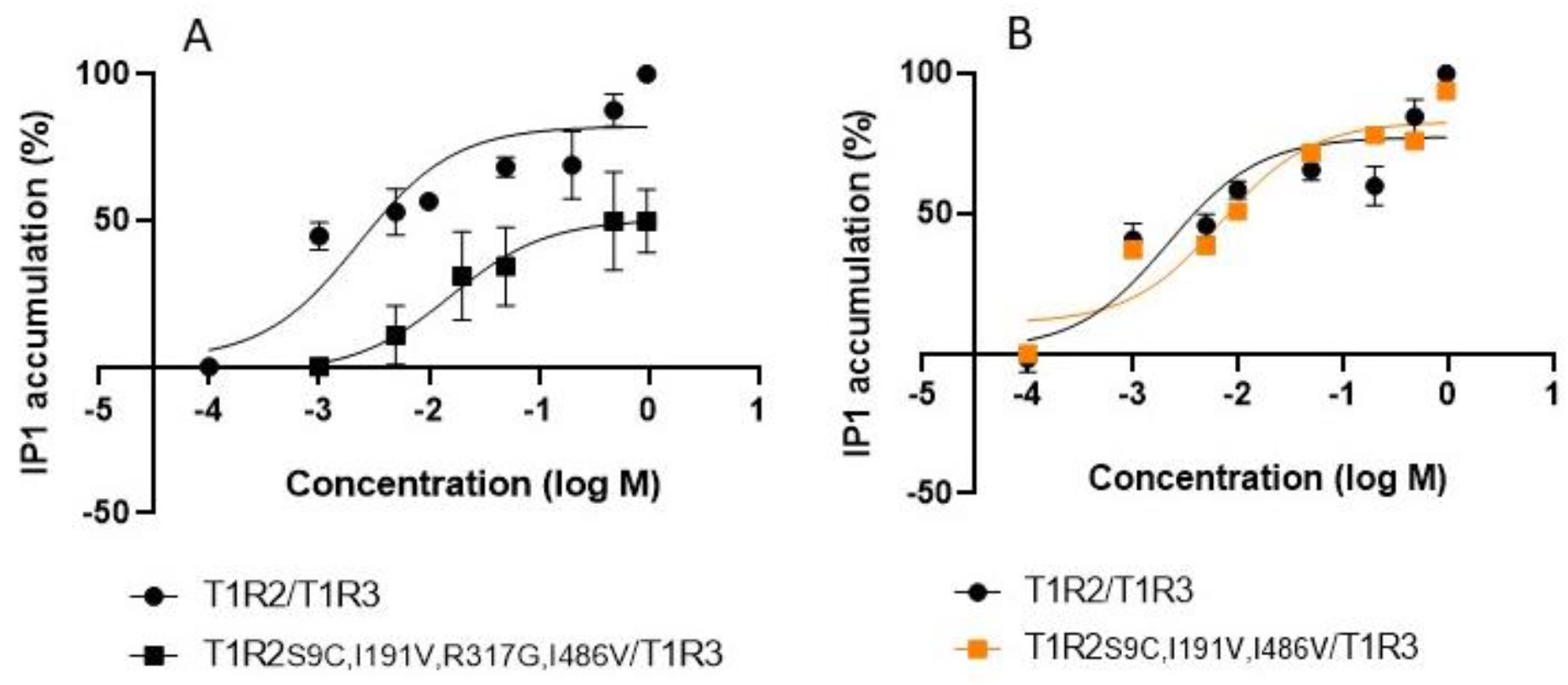
Dose-response curves of HEK293T cells transiently transfected with T1R2/T1R3 (black circle); or **(A)**T1R2S9C,I191V,R317G,I486V/T1R3 (black square); or **(B)**T1R2S9C,I191V,I486V/T1R3 (orange square), determined by using HTRF IP1-based assay, after application of L-glucose at different concentrations. WT T1R2 is mentioned as T1R2; WT T1R3 is mentioned as T1R3.

To find out which of the existing SNPs causes the reduced sensitivity, different T1R2 mutants were obtained by site directed mutagenesis. As shown in Figure 2B, in the comparison of HEK293T cells transfected with either T1R2/T1R3, or T1R2S9C,I191V,I486V/T1R3, same E_max_ value was observed (E_max_= 100%), while EC_50_ values were in the same order of magnitude (0.002 M vs. 0.006 M, respectively). Hence we suggest that the origin of difference is R317G, and not S9C, I191 or I486V.

Next, three versions of the mutated T1R2 plasmids, containing the arginine substitution to glycine in residue 317 (T1R2S9C,R317G,I486V, T1R2R317G,I486V, T1R2R317G) were examined, transfected together with T1R3, and compared to WT T1R2/T1R3. As expected, glycine instead of arginine in residue 317 increased EC_50_ value by one order of magnitude (0.015 M compared to 0.002 M, respectively), and reduced the E_max_ (∼60%) (Figure 3). These findings indicate that residue 317 is a key residue in the activation by L-glucose, and that substitution of arginine by glycine in this position negatively affects both potency and efficacy.

**Figure 3:**
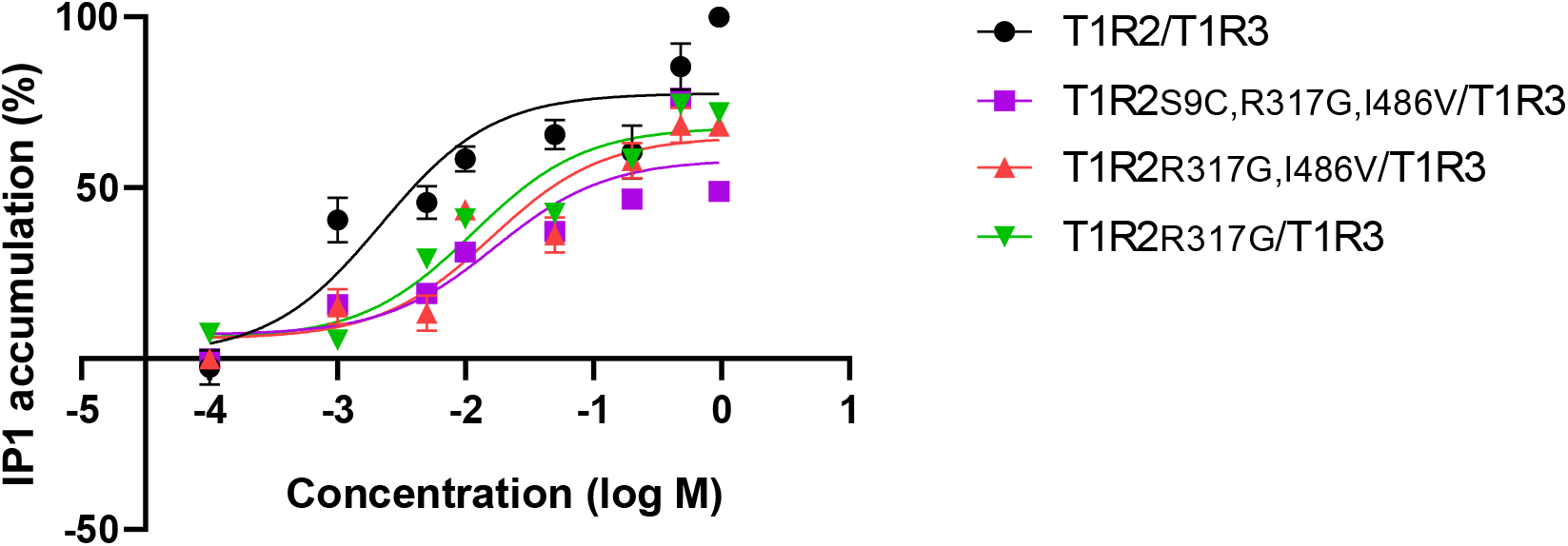
Dose-response curves of HEK293T cells transiently transfected with T1R2/T1R3 (black); T1R2S9C,R317G,I486V/T1R3 (purple); T1R2R317G,I486V/T1R3 (red); or T1R2R317G/T1R3 (green), determined by using HTRF IP1-based assay, after application of L-glucose at different concentrations. WT T1R2 is mentioned as T1R2; WT T1R3 is mentioned as T1R3.

Rreproducing our previous findings (Dubovski N, Y Ben Shoshan-Galeczki, et al., 2022), we show that both D- and L-glucose evoked activation of T1R2/T1R3 HEK293T cells in a dose response manner, also in the usage of the WT T1R2 (Figure 4). In order to determine the role of T1R2 and T1R3 in the activation by D- and L-glucose, we transformed partial transfections of either of the two components, along with Gα_16gus44_. In cells transfected with a single subunit, we assume it to function as a homodimer since class C GPCRs are obligate dimers (Kniazeff J et al., 2011). Both D- and L-glucose activated T1R3/T1R3 homodimers, with E_max_ value (90% and 100%, respectively) very close to observed in the activation of T1R2/T1R3 heterodimers (100%). T1R2/T1R2 homodimers were activated as well by D- and L-glucose, though in lower E_max_ values (80% and 50%, respectively) (Figure 4). Interestingly, different shapes of the curves were observed for D- and L-glucose: while L-glucose reaches saturation, D-glucose responds with higher IP1 levels as the concentration increased, with no saturation (Figure 4).

**Figure 4:**
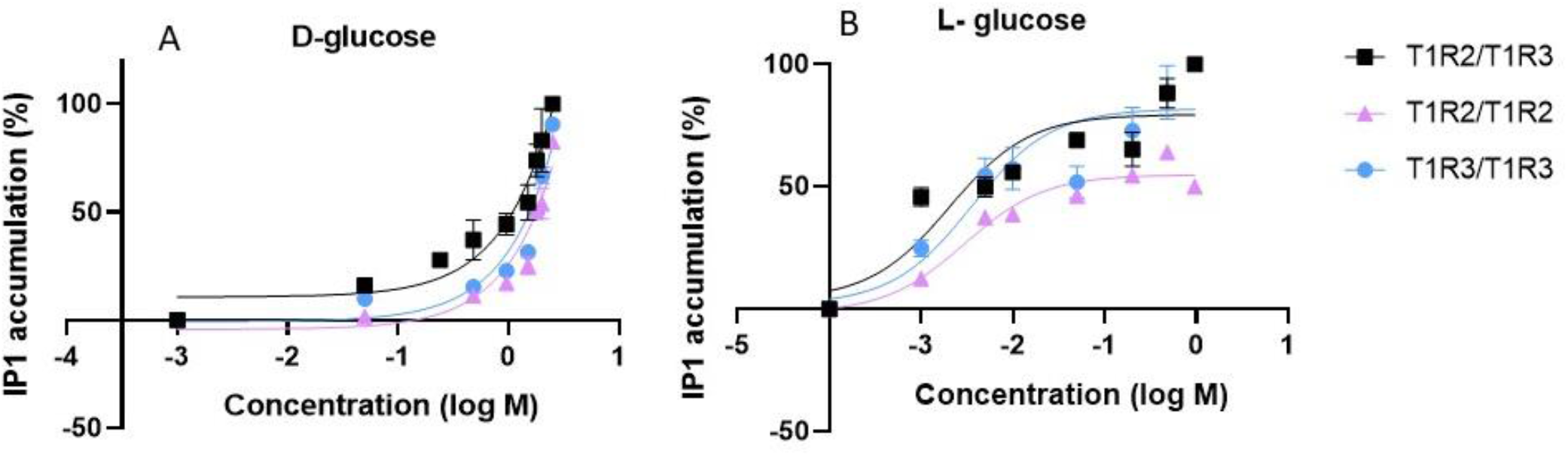
Dose-response curves of HEK293T cells transiently transfected with T1R2/T1R3 (black); or T1R2 (pink); or T1R3 (blue), determined by using Homogeneous Time-Resolved Fluorescence (HTRF) IP1-based assay, after application of **(A)** D-glucose and **(B)** L-glucose, at different concentrations. WT T1R2 is mentioned as T1R2; WT T1R3 is mentioned as T1R3.

The binding site of sugars in the VFT domain of T1R2 is well established (Dubovski N, Y Ben Shoshan-Galeczki, et al., 2022; Masuda K et al., 2012), We assumed that also in T1R3 subunit, the interaction site will be in the analogous VFT binding site.

Residue S147 in T1R3 is located in the VFT domain (Figure 5A) and plays a key role in interactions with sweeteners (Cheron JB et al., 2017). Residue S147 is well conserved among class C GPCRs (Silve C. et al., 2005). In another class C GPCR, CaSR (calcium sensing receptor), serine mutation to alanine in residue 147 was found to significantly reduce the response to calcium (Brauner-Osborne H et al., 1999; Silve Caroline et al., 2005). Computational analysis predicted that serine in residue 147 interacts with both D- and L-glucose enantiomers (Figure 5B-C). Hence we evaluated the effect of S147A in T1R3 on the response of the sweet taste receptor on D- and L-glucose.

**Figure 5:**
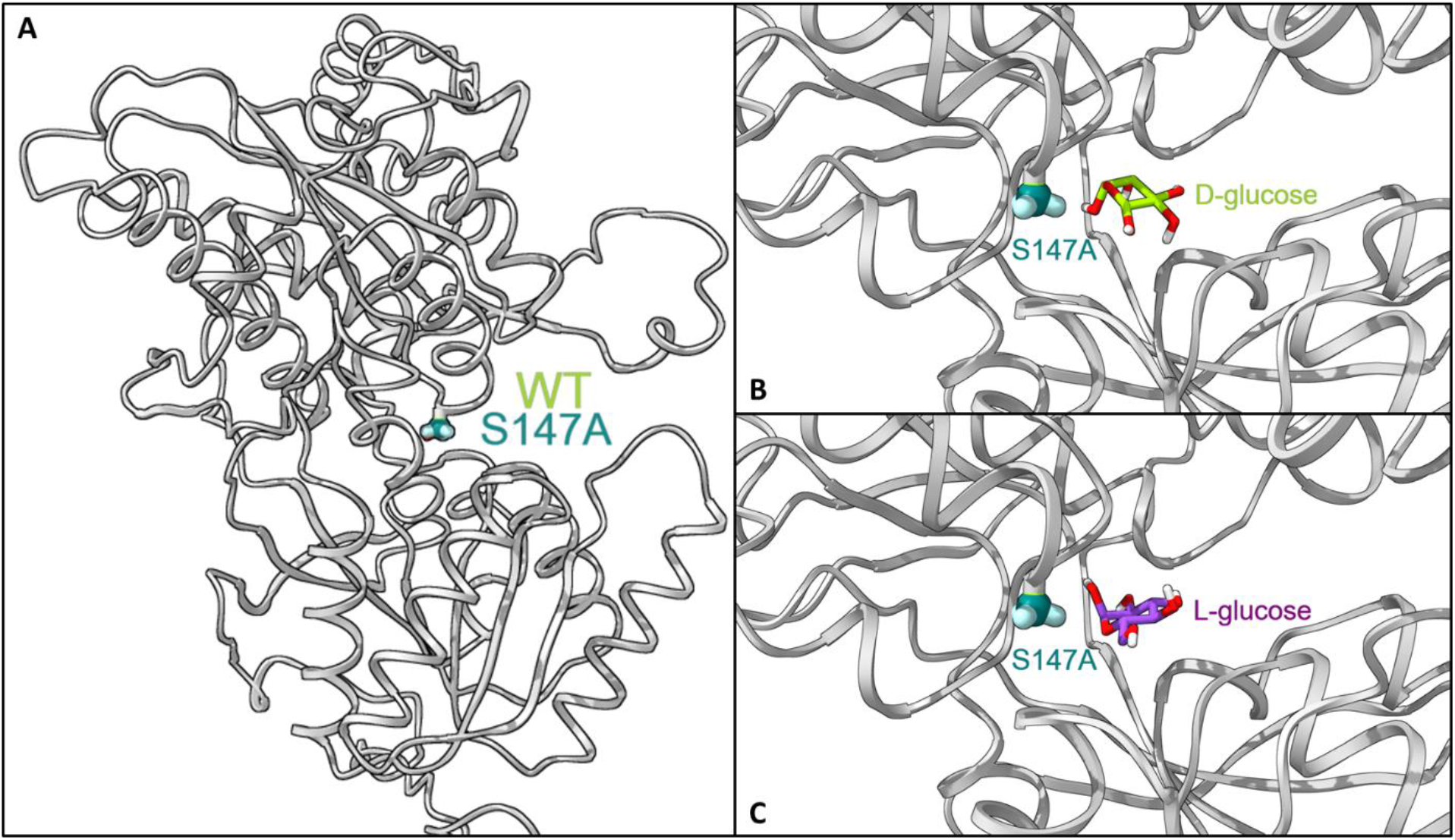
**A**. Visualization of position S147, located on the VFT domain. **B**. Docked D-glucose in the VFT domain of mutated T1R3. **C**. Docked L-glucose in the VFT domain of mutated T1R3.

As shown in Figure 6, several types of transfections were compared. Both D- and L-glucose activated T1R3/T1R3 homodimers (blue curves) with E_max_ values of 90% and 100%, respectively; and T1R2/T1R2 homodimers (purple curves) with E_max_ values of 80% and 50%, respectively; while 100% activation was observed in the case of T1R2/T1R3 (green curves). For both sugars, transfection with T1R3S147A without the presence of T1R2 (brown curves), completely abolished the activation. However, when transfected with T1R2/T1R3S147A (orange curves), cells have shown similar E_max_ level as for T1R2 only (purple curves).

**Figure 6:**
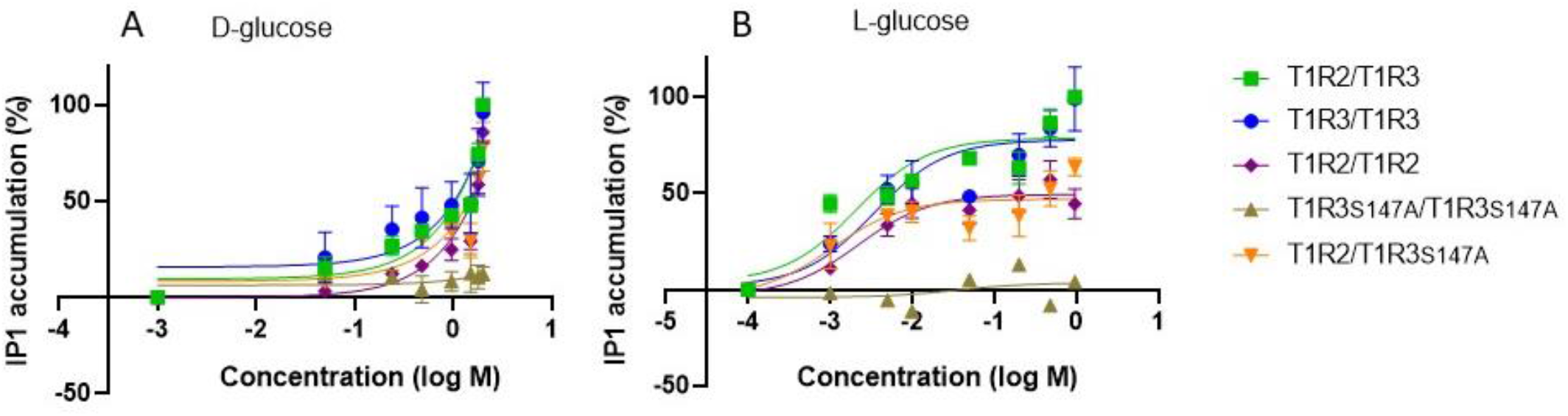
Dose-response curves of HEK293T cells transiently transfected with either T1R2/T1R3(green); or T1R3 (blue); or T1R2 (purple); or T1R3S147A (brown); or T1R2/T1R3S147A (orange), determined by using HTRF IP1-based assay, after application of **(A)** D-glucose and **(B)** L-glucose, at different concentrations. WT T1R2 is mentioned as T1R2; WT T1R3 is mentioned as T1R3.

## Materials and methods

### Materials

D-glucose, L-glucose, Dulbecco’s Modified Eagle’s medium (DMEM), lithium chloride (LiCl) and Poly-D-Lysine (PDL) were purchased from Sigma-Aldrich (St. Louis, Missouri, United States). LipofectamineTM 2000 was purchased from Invitrogen (Waltham, Massachusetts, United States). Fetal Bovine Serum (FBS) was purchased from gibco™ (Thermo Fisher Scientific). L-glutamine amino acid and penicillin–streptomycin were purchased from biological industries (Connecticut, United States).

### Cell culture

HEK293T cells, obtained from the American Type Culture Collection (ATCC), were maintained in 10% (v/v) DMEM supplemented with 10% (v/v) FBS, 1% (v/v) l-glutamine amino acid, and 0.2% (v/v) penicillin–streptomycin. Cells were kept in the incubator at 37 °C, in a humidified atmosphere containing 5% CO2.

### IP1 assay

The protocol is described in detail in our previous work (Dubovski N, Y Ben Shoshan-Galeczki, et al., 2022). Transfections were performed on a 10-cm plate. The desired plasmids were transiently transfected into HEK293T cells using LipofectamineTM 2000. In addition to the plasmids described in the Results, all transfections were performed with another plasmid (pcDNA3) containing Gα_16gus44_, even if not mentioned. Transfected cells were seeded onto a 24-well culture plate coated with PDL. On the experiment day, cells were exposed to the ligands dissolved in 0.05 M LiCl, because LiCl is crucial for IP1 accumulation. Cells were exposed to the tastants by adding the tastant solution directly into the wells, for 5 min. At the end of the exposure time, the well content was replaced with fresh 0.1% DMEM containing LiCl, for another 55-min incubation. The homogeneous time-resolved fluorescence IP1 assay was used to measure intracellular IP1 accumulation according to the Cisbio IP-One Gq kit (Cisbio Bioassays, Perkin Elmer). Dose-response curves were fitted by Prism 8 (Graphpad). Each of the graphs represents at least 3 independent data sets.

### Sequencing

Sequencing of T1R2 was performed by Sanger dideoxy method (Zimmermann J et al., 1988).

### Site directed mutagenesis

Site directed mutagenesis in T1R2 and T1R3 plasmids were performed by Transfer-PCR (TPCR), as described previously (Erijman A et al., 2011).

### Modeling and docking

Computational analysis was described previously (Dubovski N, Y Ben Shoshan-Galeczki, et al., 2022). In brief, D- and L-glucose were downloaded from PubChem, and were prepared for docking using LigPrep (Schrodinger tools 2017–2). Conformers, tautomers and protomers (different protonation states of ligands) were enumerated at pH 7.0 ± 0.5, retaining specified chiral centers (Maestro Version 2021-1). The T1R3 model was downloaded from UniProt AlphaFold database (https://alphafold.ebi.ac.uk/entry/Q7RTX0, version 1) and was prepared with Schrodinger Maestro (2021-1). The binding site was defined as 12 angstrom around the docked ligand (glutamine) from the fish X-ray PDB structure (5×2M, chain B). The docking parameters were set to Glide XP flexible docking, with extended sampling.

### Interactions analysis

Interactions analysis was performed with the sum of interactions fingerprints of heavy atoms within 4Å from docked ligands as described for analysis of sub-pockets (Deng Z et al., 2004). The interactions between the ligands and the residues within the binding site were analyzed with Schrodinger Interactions Fingerprints tool (2021-3), with the default definitions. We calculated the total interactions for each residue different docked poses of α- D-glucose, β- D-glucose, α- L-glucose and β- L-glucose and divided the sum of per-residue interactions by the number of total docked poses.

### Multiple sequence alignment

Amino acid sequences of T1R2, T1R3, CaSR, and metabotropic glutamate receptors 1-8 (mGluRs) were obtained from Pubmed Protein (https://www.ncbi.nlm.nih.gov/protein/). Reference sequences of T1R2 were obtained from Pubmed Nucleotide (https://www.ncbi.nlm.nih.gov/nucleotide/). Alignments were performed with the Clustal Omega server (https://www.ebi.ac.uk/Tools/msa/clustalo/).

## Discussion

This study is a continuation of our previous work, where we showed that the sweetness of both L and D-glucose enantiomers is mediated via T1R2/T1R3 (Dubovski N, Y Ben Shoshan-Galeczki, et al., 2022). In the current study we aimed to compare two T1R2 sequences for their sensitivity in L-glucose activation: the original RefSeq (NM_152232.1), and the updated RefSeq (NM_152232.4). The original RefSeq, which have been used in our previous study, contains 4 SNPs compared to NM_152232.4, in positions 9, 191, 317 and 486. Then, using NM_152232.4, we aimed to reproduce our previous finding regarding D- and L-glucose activation via T1R2/T1R3, and to understand the contribution of each of the sweet receptor subunits in D- and L-glucose recognition. We also aimed to confirm our hypothesis that it is the VFT domain of T1R3 that is involved in recognizing D- and L-glucose.

In T1R2, 9 SNPs with frequency greater than 1% exist among the human world population (Smith NJ et al., 2021). Four of these SNPs differ between the updated sequence (NM_152232.4) and the previous sequence (NM_152232.1). Among these SNPs, there is the substitution of arginine with glycine in residue 317 (MAF=0.238) (Smith NJ et al., 2021). Here we found that these “reference sequences” have significantly different sensitivities, and tracked the difference down to the SNP R317G. This position (not conserved among Class C GPCRs) is responsible to the increased sensitivity of the “new WT” sequence. The substitution of arginine to glycine in residue 317, which is located outside of the binding site, might affect trafficking to the cell membrane, receptor stability or have allosteric effect on binding. While our studies did not discern the mechanisms of the effect, the established effect on sensitivity of the receptor has important implications both for cell-based studies and for genotyping populations. Position 317 is not conserved among Class C GPCRs (Figure S1). T1R3 has alanine at position 317 and its potential role in T1R3 sensitivity to ligands requires further study. Since Feb 12 2020, two updated reference sequences were released: NM_152232.5, and NM_152232.6. However, the amino acid sequence in the coding region of NM_152232.4 remained the same (Figure S2).

Next we studied the roles of the T1R subunits. The canonical taste receptor is a heterodimer of T1R2 and T1R3. The sweet taste receptor is expressed not only in the oral cavity, but also in extra-oral tissues (Dubovski N, F Fierro, et al., 2022), where additionally to T1R2/T1R3, it can also function as a homodimer of T1R3/T1R3 (Buchanan KL et al., 2022; Li X, 2009; Margolskee RF et al., 2007; Ohtsu Y et al., 2014; Zhao GQ et al., 2003). Here for the first time we show that D- and L-glucose can activate T1R2/T1R2 and T1R3/T1R3 homodimers, with different levels of activation. Our findings are in accord with other studies (Li X, 2009; Servant G et al., 2020) which have shown responses to allosteric ligands and to high sucrose levels, in the transfection with either T1R2 or T1R3, respectively.

To confirm the binding region for D- and L-glucose in T1R3, we carried out mutagenesis of a key residue, 147. This position is well conserved among Class C GPCRs, including T1R3 (Figure S1). This residue is part of T1R3 binding site, and was shown to be involved in the binding of sweeteners (Cheron JB et al., 2017). In CaSR, another Class C GPCR, serine substitution to alanine in residue 147 reduced the responses to calcium (Brauner-Osborne H et al., 1999; Silve C. et al., 2005). Our computational analysis predicted S147 to interact with each of the glucose enantiomers. Indeed, mutation of serine to alanine in residue 147 in T1R3 abolished T1R3S147A-transfected cells activation by both D- and L-glucose. However, in the transfection of T1R3S147A with T1R2, similar E_max_ value was observed, as in T1R2 transfection alone.

An important finding is the distinguishable and characteristic curves of D- and L-glucose. Although the examined genetic variations affect both enantiomer similarly, the shape of the dose response curves is different: while L-glucose reaches an early plateau, no saturation was observed for D-glucose. Interestingly, the typical shapes for D- and L-glucose are conserved also for partial transfections. This characteristic of D-glucose, of not reaching saturation, was also shown in sensory studies (Low JY et al., 2017). The reason to this distinct curve shapes may be related to the sugar characteristics (caloric vs. non-caloric) and requires further study.

## Conclusion

As the reference sequences keep changing, it is critical to indicate the RefSeq number in sweet taste receptor studies, as suggested by (Smith NJ et al., 2021). We show that RefSeq T1R2 NM_152232.4 is more sensitive than the previous one (NM_152232.1) in cell-based functional assays. We found that it is the naturally occurring glycine at position 317 in T1R2 that is responsible for reduced receptor sensitivity of NM_152232.1. D-glucose and its enantiomeric non-calorie sugar, L-glucose, can activate T1R2 and T1R3 transfected without the partner subunit in dose response but with distinctly different curve shapes. Mutation of S147 to alanine in T1R3 abolished activation by glucose, but transfection of T1R3S147A with T1R2 led to same E_max_ as T1R2/T1R2.

## Acknowledgments

The authors thank Dr. R. F. Margolskee for the pcDNA3 of chimeric Gα_16gust44_ and Dr. Maik Behrens for the pcDNA3 of T1R3, and for the pcDNA5FRT PM of T1R2S9C,I191V,R317G,I486V. We thank Dr. Yoav Peleg for help with sequencing and site directed mutagenesis; and for the construction of pcDNA5 of T1R2, pcDNA5 of T1R2S9C,I191V,I486V, pcDNA5 of T1R2S9C,R317G,I486V, pcDNA5 of T1R2R317G,I486V, pcDNA5 of T1R2R317G, and pcDNA3 of TIR3S147A. The study was supported by ISF grant #1129/19.

## References

Archer, N. S., Liu, D., Shaw, J., Hannan, G., Duesing, K., & Keast, R. (2016). A Comparison of Collection Techniques for Gene Expression Analysis of Human Oral Taste Tissue. PLoS One, 11(3), e0152157. doi: 10.1371/journal.pone.0152157

Brauner-Osborne, H., Jensen, A. A., Sheppard, P. O., O’Hara, P., & Krogsgaard-Larsen, P. (1999). The agonist-binding domain of the calcium-sensing receptor is located at the amino-terminal domain. J Biol Chem, 274(26), 18382–18386. doi: 10.1074/jbc.274.26.18382

Buchanan, K. L., Rupprecht, L. E., Kaelberer, M. M., Sahasrabudhe, A., Klein, M. E., Villalobos, J. A., Liu, W. W., Yang, A., Gelman, J., Park, S., Anikeeva, P., & Bohorquez, D. V. (2022). The preference for sugar over sweetener depends on a gut sensor cell. Nat Neurosci, 25(2), 191–200. doi: 10.1038/s41593-021-00982-7

Chamoun, E., Mutch, D. M., Allen-Vercoe, E., Buchholz, A. C., Duncan, A. M., Spriet, L. L., Haines, J., Ma, D. W. L., & Guelph Family Health, S. (2018). A review of the associations between single nucleotide polymorphisms in taste receptors, eating behaviors, and health. Crit Rev Food Sci Nutr, 58(2), 194–207. doi: 10.1080/10408398.2016.1152229

Cheron, J. B., Golebiowski, J., Antonczak, S., & Fiorucci, S. (2017). The anatomy of mammalian sweet taste receptors. Proteins, 85(2), 332–341. doi: 10.1002/prot.25228

Deng, Z., Chuaqui, C., & Singh, J. (2004). Structural interaction fingerprint (SIFt): a novel method for analyzing three-dimensional protein-ligand binding interactions. J Med Chem, 47(2), 337–344. doi: 10.1021/jm030331x

DuBois, G. E. (2016). Molecular mechanism of sweetness sensation. Physiol Behav, 164(Pt B), 453–463. doi: 10.1016/j.physbeh.2016.03.015

Dubovski, N., Ben Shoshan-Galeczki, Y., Malach, E., & Niv, M. Y. (2022). Taste and chirality: l-glucose sweetness is mediated by TAS1R2/TAS2R3 receptor. Food Chem, 373(Pt A), 131393. doi: 10.1016/j.foodchem.2021.131393

Dubovski, N., Fierro, F., Margulis, E., Ben Shoshan-Galeczki, Y., Peri, L., & Niv, M. Y. (2022). Taste GPCRs and their ligands. Prog Mol Biol Transl Sci, 193(1), 177–193. doi: 10.1016/bs.pmbts.2022.06.008

Eny, K. M., Wolever, T. M., Corey, P. N., & El-Sohemy, A. (2010). Genetic variation in TAS1R2 (Ile191Val) is associated with consumption of sugars in overweight and obese individuals in 2 distinct populations. Am J Clin Nutr, 92(6), 1501–1510. doi: 10.3945/ajcn.2010.29836

Erijman, A., Dantes, A., Bernheim, R., Shifman, J. M., & Peleg, Y. (2011). Transfer-PCR (TPCR): a highway for DNA cloning and protein engineering. J Struct Biol, 175(2), 171–177. doi: 10.1016/j.jsb.2011.04.005

Fushan, A. A., Simons, C. T., Slack, J. P., Manichaikul, A., & Drayna, D. (2009). Allelic polymorphism within the TAS1R3 promoter is associated with human taste sensitivity to sucrose. Curr Biol, 19(15), 1288–1293. doi: 10.1016/j.cub.2009.06.015

Haznedaroglu, E., Koldemir-Gunduz, M., Bakir-Coskun, N., Bozkus, H. M., Cagatay, P., Susleyici-Duman, B., & Mentes, A. (2015). Association of sweet taste receptor gene polymorphisms with dental caries experience in school children. Caries Res, 49(3), 275–281. doi: 10.1159/000381426

Jang, H. J., Kokrashvili, Z., Theodorakis, M. J., Carlson, O. D., Kim, B. J., Zhou, J., Kim, H. H., Xu, X., Chan, S. L., Juhaszova, M., Bernier, M., Mosinger, B., Margolskee, R. F., & Egan, J. M. (2007). Gut-expressed gustducin and taste receptors regulate secretion of glucagon-like peptide-1. Proc Natl Acad Sci U S A, 104(38), 15069–15074. doi: 10.1073/pnas.0706890104

Jiang, P., Cui, M., Zhao, B., Liu, Z., Snyder, L. A., Benard, L. M., Osman, R., Margolskee, R. F., & Max, M. (2005). Lactisole interacts with the transmembrane domains of human T1R3 to inhibit sweet taste. J Biol Chem, 280(15), 15238–15246. doi: 10.1074/jbc.M414287200

Kniazeff, J., Prezeau, L., Rondard, P., Pin, J. P., & Goudet, C. (2011). Dimers and beyond: The functional puzzles of class C GPCRs. Pharmacol Ther, 130(1), 9–25. doi: 10.1016/j.pharmthera.2011.01.006

Kulkarni, G. V., Chng, T., Eny, K. M., Nielsen, D., Wessman, C., & El-Sohemy, A. (2013). Association of GLUT2 and TAS1R2 genotypes with risk for dental caries. Caries Res, 47(3), 219–225. doi: 10.1159/000345652

Levin, G. V., Zehner, L. R., Saunders, J. P., & Beadle, J. R. (1995). Sugar substitutes: their energy values, bulk characteristics, and potential health benefits. Am J Clin Nutr, 62(5 Suppl), 1161S–1168S. doi: 10.1093/ajcn/62.5.1161S

Li, X. (2009). T1R receptors mediate mammalian sweet and umami taste. Am J Clin Nutr, 90(3), 733S–737S. doi: 10.3945/ajcn.2009.27462G

Li, X., Staszewski, L., Xu, H., Durick, K., Zoller, M., & Adler, E. (2002). Human receptors for sweet and umami taste. Proc Natl Acad Sci U S A, 99(7), 4692–4696. doi: 10.1073/pnas.072090199

Low, J. Y., McBride, R. L., Lacy, K. E., & Keast, R. S. (2017). Psychophysical Evaluation of Sweetness Functions Across Multiple Sweeteners. Chem Senses, 42(2), 111–120. doi: 10.1093/chemse/bjw109

Margolskee, R. F., Dyer, J., Kokrashvili, Z., Salmon, K. S., Ilegems, E., Daly, K., Maillet, E. L., Ninomiya, Y., Mosinger, B., & Shirazi-Beechey, S. P. (2007). T1R3 and gustducin in gut sense sugars to regulate expression of Na+-glucose cotransporter 1. Proc Natl Acad Sci U S A, 104(38), 15075–15080. doi: 10.1073/pnas.0706678104

Masuda, K., Koizumi, A., Nakajima, K., Tanaka, T., Abe, K., Misaka, T., & Ishiguro, M. (2012). Characterization of the modes of binding between human sweet taste receptor and low-molecular-weight sweet compounds. PLoS One, 7(4), e35380. doi: 10.1371/journal.pone.0035380

Nelson, G., Hoon, M. A., Chandrashekar, J., Zhang, Y., Ryba, N. J., & Zuker, C. S. (2001). Mammalian sweet taste receptors. Cell, 106(3), 381–390. doi: 10.1016/s0092-8674(01)00451-2

Nie, Y., Vigues, S., Hobbs, J. R., Conn, G. L., & Munger, S. D. (2005). Distinct contributions of T1R2 and T1R3 taste receptor subunits to the detection of sweet stimuli. Curr Biol, 15(21), 1948–1952. doi: 10.1016/j.cub.2005.09.037

Ohtsu, Y., Nakagawa, Y., Nagasawa, M., Takeda, S., Arakawa, H., & Kojima, I. (2014). Diverse signaling systems activated by the sweet taste receptor in human GLP-1-secreting cells. Mol Cell Endocrinol, 394(1-2), 70–79. doi: 10.1016/j.mce.2014.07.004

Pioltine, M. B., de Melo, M. E., Santos, A. S., Machado, A. D., Fernandes, A. E., Fujiwara, C. T., Cercato, C., & Mancini, M. C. (2018). Genetic Variations in Sweet Taste Receptor Gene Are Related to Chocolate Powder and Dietary Fiber Intake in Obese Children and Adolescents. J Pers Med, 8(1). doi: 10.3390/jpm8010007

Serrano, J., Seflova, J., Park, J., Pribadi, M., Sanematsu, K., Shigemura, N., Serna, V., Yi, F., Mari, A., Procko, E., Pratley, R. E., Robia, S. L., & Kyriazis, G. A. (2021). The Ile191Val is a partial loss-of-function variant of the TAS1R2 sweet-taste receptor and is associated with reduced glucose excursions in humans. Mol Metab, 54, 101339. doi: 10.1016/j.molmet.2021.101339

Servant, G., Kenakin, T., Zhang, L., Williams, M., & Servant, N. (2020). The function and allosteric control of the human sweet taste receptor. Adv Pharmacol, 88, 59–82. doi: 10.1016/bs.apha.2020.01.002

Silve, C., Petrel, C., Leroy, C., Bruel, H., Mallet, E., Rognan, D., & Ruat, M. (2005). Delineating a Ca2+ binding pocket within the venus flytrap module of the human calcium-sensing receptor. J Biol Chem, 280(45), 37917–37923. doi: 10.1074/jbc.M506263200

Silve, C., Petrel, C., Leroy, C., Bruel, H., Mallet, E., Rognan, D., & Ruat, M. (2005). Delineating a Ca2+ Binding Pocket within the Venus Flytrap Module of the Human Calcium-sensing Receptor. Journal of Biological Chemistry, 280(45), 37917–37923. doi: 10.1074/jbc.M506263200

Smith, N. J., Grant, J. N., Moon, J. I., So, S. S., & Finch, A. M. (2021). Critically evaluating sweet taste receptor expression and signaling through a molecular pharmacology lens. FEBS J, 288(8), 2660–2672. doi: 10.1111/febs.15768

Treesukosol, Y., Smith, K. R., & Spector, A. C. (2011). The functional role of the T1R family of receptors in sweet taste and feeding. Physiol Behav, 105(1), 14–26. doi: 10.1016/j.physbeh.2011.02.030

Winnig, M., Bufe, B., Kratochwil, N. A., Slack, J. P., & Meyerhof, W. (2007). The binding site for neohesperidin dihydrochalcone at the human sweet taste receptor. BMC Struct Biol, 7, 66. doi: 10.1186/1472-6807-7-66

Xi, R., Zheng, X., & Tizzano, M. (2022). Role of Taste Receptors in Innate Immunity and Oral Health. J Dent Res, 101(7), 759–768. doi: 10.1177/00220345221077989

Xu, H., Staszewski, L., Tang, H., Adler, E., Zoller, M., & Li, X. (2004). Different functional roles of T1R subunits in the heteromeric taste receptors. Proc Natl Acad Sci U S A, 101(39), 14258–14263. doi: 10.1073/pnas.0404384101

Zhao, G. Q., Zhang, Y., Hoon, M. A., Chandrashekar, J., Erlenbach, I., Ryba, N. J., & Zuker, C. S. (2003). The receptors for mammalian sweet and umami taste. Cell, 115(3), 255–266. doi: 10.1016/s0092-8674(03)00844-4

Zimmermann, J., Voss, H., Schwager, C., Stegemann, J., & Ansorge, W. (1988). Automated Sanger dideoxy sequencing reaction protocol. FEBS Lett, 233(2), 432–436. doi: 10.1016/0014-5793(88)80477-0

