## Supplementary material for "Sensitivity of human sweet taste receptor subunits T1R2 and T1R3 to activation by glucose enantiomers": Figure S1, Figure S2

### Supplementary data

```
CaSR            FDTCNTVSKALEATLSFVAQNKIDSLNL-DEFCNCSEHIP---------STIAVVGATGS        147
mGluR1          RDSCWHSSVALEQSIEFIRDSLISIRDEKDGINRCLPDGQSLPPGRTKKPIAGVIGPGSS        165
mGluR5          RDSCWHSAVALEQSIEFIRDSLISSE-EEEGLVRCVDGSS--SSFRSKKPIVGVIGPGSS        152
mGluR6          LDTCSRDTYALEQALSFVQALIRGRGDGDEVGVRCPGGVPPLRP-APPERVVAVVGASAS        154
mGluR7          LDTCSRDTYALEQSLTFVQALIQKD----TSDVRCTNGEPPVFV--KPEKVVGVIGASGS        159
mGluR4          LDTCSRDTHALEQSLTFVQALIEKD----GTEVRCGSGGPPIIT--KPERVVGVIGASGS        159
mGluR8          LDTCSRDTYALEQSLTFVQALIEKD----ASDVKCANGDPPIFT--KPDKISGVIGAAAS        156
mGluR2          LDSCSKDTHALEQALDFVRASLSRGAD--GSRHICPDGSYATHG-DAPTAITGVIGGSYS        145
mGluR3          LDTCSRDTYALEQSLEFVRASLTKV-D--EAEYMCPDGSYAIQE-NIPLLIAGVIGGSYS        151
T1R2            VDVCYISN-NVQPVLYFLAHE-DNLL---PIQEDYSNY---------ISRVVAVIGPDNS        144
T1R3            FDTCSEPVVAMKPSLMFLAKAGSRDI---AAYCNYTQY---------QPRVLAVIGPHSS        147
                 * *      ::  : *:                                  .*:*   *

CaSR            GVSTAVANLLGLFYIPQVSYASSSRLLSNKNQFKSFLRTIPNDEHQATAMADIIEYFRWN        207
mGluR1          SVAIQVQNLLQLFDIPQIAYSATSIDLSDKTLYKYFLRVVPSDTLQARAMLDIVKRYNWT        225
mGluR5          SVAIQVQNLLQLFNIPQIAYSATSMDLSDKTLFKYFMRVVPSDAQQARAMVDIVKRYNWT        212
mGluR6          SVSIMVANVLRLFAIPQISYASTAPELSDSTRYDFFSRVVPPDSYQAQAMVDIVRALGWN        214
mGluR7          SVSIMVANILRLFQIPQISYASTAPELSDDRRYDFFSRVVPPDSFQAQAMVDIVKALGWN        219
mGluR4          SVSIMVANILRLFKIPQISYASTAPDLSDNSRYDFFSRVVPSDTYQAQAMVDIVRALKWN        219
mGluR8          SVSIMVANILRLFKIPQISYASTAPELSDNTRYDFFSRVVPPDSYQAQAMVDIVTALGWN        216
mGluR2          DVSIQVANLLRLFQIPQISYASTSAKLSDKSRYDYFARTVPPDFFQAKAMAEILRFFNWT        205
mGluR3          SVSIQVANLLRLFQIPQISYASTSAKLSDKSRYDYFARTVPPDFYQAKAMAEILRFFNWT        211
T1R2            ESVMTVANFLSLFLLPQITYSAISDELRDKVRFPALLRTTPSADHHIEAMVQLMLHFRWN        204
T1R3            ELAMVTGKFFSFFLMPQVSYGASMELLSARETFPSFFRTVPSDRVQLTAAAELLQEFGWN        207
                     . :.: :* :**::*.:    *     :  : *. *    :  *  :::    *.

CaSR            WVGTIAADDDYGRPGIEKFREEAE-ERDICIDFSELISQYSD--------EEEIQHVVEV        258
mGluR1          YVSAVHTEGNYGESGMDAFKELAA-QEGLCIAHSDKIYSNAG--------EKSFDRLLRK        276
mGluR5          YVSAVHTEGNYGESGMEAFKDMSA-KEGICIAHSYKIYSNAG--------EQSFDKLLKK        263
mGluR6          YVSTLASEGNYGESGVEAFVQISREAGGVCIAQSIKIPREPK--------PGEFSKVIRR        266
mGluR7          YVSTLASEGSYGEKGVESFTQISKEAGGLCIAQSVRIPQERKDR------TIDFDRIIKQ        273
mGluR4          YVSTVASEGSYGESGVEAFIQKSREDGGVCIAQSVKIPREPK--------AGEFDKIIRR        271
mGluR8          YVSTLASEGNYGESGVEAFTQISREIGGVCIAQSQKIPREPR--------PGEFEKIIKR        268
mGluR2          YVSTVASEGDYGETGIEAFELEAR-ARNICVATSEKVGRAMS--------RAAFEGVVRA        256
mGluR3          YVSTVASEGDYGETGIEAFEQEAR-LRNICIATAEKVGRSNI--------RKSYDSVIRE        262
T1R2            WIIVLVSSDTYGRDNGQLLGERVA-RRDICIAFQETLPTLQPNQNMTSEERQRLVTIVDK        263
T1R3            WVAALGSDDEYGRQGLSIFSALAA-ARGICIAHEGLVPLPRADDSR----LGKVQDVLHQ        262
                :: .: :.. **. . . :        .:*:     :                   ::

CaSR            I-Q-NSTAKVIVVFSSGPDLEPLIKEIVRRNITGK-IWLASEAWASSSLIAMPQYFHVVG        315
mGluR1          LRERLPKARVVVCFCEGMTVRGLLSAMRRLGVVGEFSLIGSDGWADRDEV-IEGYEVEAN        335
mGluR5          LTSHLPKARVVACFCEGMTVRGLLMAMRRLGLAGEFLLLGSDGWADRYDV-TDGYQREAV        322
mGluR6          LME-TPNARGIIIFANEDDIRRVLEAARQANLTGHFLWVGSDSWGAKTSP-ILSLEDVAV        324
mGluR7          LLD-TPNSRAVVIFANDEDIKQILAAAKRADQVGHFLWVGSDSWGSKINP-LHQHEDIAE        331
mGluR4          LLE-TSNARAVIIFANEDDIRRVLEAARRANQTGHFFWMGSDSWGSKIAP-VLHLEEVAE        329
mGluR8          LLE-TPNARAVIMFANEDDIRRILEAAKKLNQSGHFLWIGSDSWGSKIAP-VYQQEEIAE        326
mGluR2          LLQ-KPSARVAVLFTRSEDARELLAASQRLNA--SFTWVASDGWGALESV-VAGSEGAAE        312
mGluR3          LLQ-KPNARVVVLFMRSDDSRELIAAASRANA--SFTWVASDGWGAQESI-IKGSEHVAY        318
T1R2            LQ--QSTARVVVVFSPDLTLYHFFNEVLRQNFTGA-VWIASESWAIDPVLHNLTELRHLG        320
T1R3            VN--QSSVQVVLLFASVHAAHALFNYSISSRLSPK-VWVASEAWLTSDLVMGLPGMAQMG        319
                :     . :    *        .:              :.*:.*
```

**Figure S1:** Sequence alignment of human T1R2, T1R3, CaSR, and metabotropic glutamate receptors 1-8 (mGluRs). Marked in red are the equivalent positions of S147 in T1R3, and R317 in T1R2.

```
NM_152232.1      MGPRAKTICSLFFLLWVLAEPAENSDFYLPGDYLLGGLFSLHANMKGIVHLNFLQVPMCK      60
NM_152232.4      MGPRAKTISSLFFLLWVLAEPAENSDFYLPGDYLLGGLFSLHANMKGIVHLNFLQVPMCK      60
NM_152232.5      MGPRAKTISSLFFLLWVLAEPAENSDFYLPGDYLLGGLFSLHANMKGIVHLNFLQVPMCK      60
NM_152232.6      MGPRAKTISSLFFLLWVLAEPAENSDFYLPGDYLLGGLFSLHANMKGIVHLNFLQVPMCK      60
                 ********.***************************************************

NM_152232.1      EYEVKVIGYNLMQAMRFAVEEINNDSSLLPGVLLGYEIVDVCYISNNVQPVLYFLAHEDN      120
NM_152232.4      EYEVKVIGYNLMQAMRFAVEEINNDSSLLPGVLLGYEIVDVCYISNNVQPVLYFLAHEDN      120
NM_152232.5      EYEVKVIGYNLMQAMRFAVEEINNDSSLLPGVLLGYEIVDVCYISNNVQPVLYFLAHEDN      120
NM_152232.6      EYEVKVIGYNLMQAMRFAVEEINNDSSLLPGVLLGYEIVDVCYISNNVQPVLYFLAHEDN      120
                 ************************************************************

NM_152232.1      LLPIQEDYSNYISRVVAVIGPDNSESVMTVANFLSLFLLPQITYSAISDELRDKVRFPAL      180
NM_152232.4      LLPIQEDYSNYISRVVAVIGPDNSESVMTVANFLSLFLLPQITYSAISDELRDKVRFPAL      180
NM_152232.5      LLPIQEDYSNYISRVVAVIGPDNSESVMTVANFLSLFLLPQITYSAISDELRDKVRFPAL      180
NM_152232.6      LLPIQEDYSNYISRVVAVIGPDNSESVMTVANFLSLFLLPQITYSAISDELRDKVRFPAL      180
                 ************************************************************

NM_152232.1      LRTTPSADHHVEAMVQLMLHFRWNWIIVLVSSDTYGRDNGQLLGERVARRDICIAFQETL      240
NM_152232.4      LRTTPSADHHIEAMVQLMLHFRWNWIIVLVSSDTYGRDNGQLLGERVARRDICIAFQETL      240
NM_152232.5      LRTTPSADHHIEAMVQLMLHFRWNWIIVLVSSDTYGRDNGQLLGERVARRDICIAFQETL      240
NM_152232.6      LRTTPSADHHIEAMVQLMLHFRWNWIIVLVSSDTYGRDNGQLLGERVARRDICIAFQETL      240
                 **********:*************************************************

NM_152232.1      PTLQPNQNMTSEERQRLVTIVDKLQQSTARVVVVFSPDLTLYHFFNEVLRQNFTGAVWIA      300
NM_152232.4      PTLQPNQNMTSEERQRLVTIVDKLQQSTARVVVVFSPDLTLYHFFNEVLRQNFTGAVWIA      300
NM_152232.5      PTLQPNQNMTSEERQRLVTIVDKLQQSTARVVVVFSPDLTLYHFFNEVLRQNFTGAVWIA      300
NM_152232.6      PTLQPNQNMTSEERQRLVTIVDKLQQSTARVVVVFSPDLTLYHFFNEVLRQNFTGAVWIA      300
                 ************************************************************

NM_152232.1      SESWAIDPVLHNLTELGHLGTFLGITIQSVPIPGFSEFREWGPQAGPPPLSRTSQSYTCN      360
NM_152232.4      SESWAIDPVLHNLTELRHLGTFLGITIQSVPIPGFSEFREWGPQAGPPPLSRTSQSYTCN      360
NM_152232.5      SESWAIDPVLHNLTELRHLGTFLGITIQSVPIPGFSEFREWGPQAGPPPLSRTSQSYTCN      360
NM_152232.6      SESWAIDPVLHNLTELRHLGTFLGITIQSVPIPGFSEFREWGPQAGPPPLSRTSQSYTCN      360
                 **************** *******************************************

NM_152232.1      QECDNCLNATLSFNTILRLSGERVVYSVYSAVYAVAHALHSLLGCDKSTCTKRVVYPWQL      420
NM_152232.4      QECDNCLNATLSFNTILRLSGERVVYSVYSAVYAVAHALHSLLGCDKSTCTKRVVYPWQL      420
NM_152232.5      QECDNCLNATLSFNTILRLSGERVVYSVYSAVYAVAHALHSLLGCDKSTCTKRVVYPWQL      420
NM_152232.6      QECDNCLNATLSFNTILRLSGERVVYSVYSAVYAVAHALHSLLGCDKSTCTKRVVYPWQL      420
                 ************************************************************

NM_152232.1      LEEIWKVNFTLLDHQIFFDPQGDVALHLEIVQWQWDRSQNPFQSVASYYPLQRQLKNIQD      480
NM_152232.4      LEEIWKVNFTLLDHQIFFDPQGDVALHLEIVQWQWDRSQNPFQSVASYYPLQRQLKNIQD      480
NM_152232.5      LEEIWKVNFTLLDHQIFFDPQGDVALHLEIVQWQWDRSQNPFQSVASYYPLQRQLKNIQD      480
NM_152232.6      LEEIWKVNFTLLDHQIFFDPQGDVALHLEIVQWQWDRSQNPFQSVASYYPLQRQLKNIQD      480
                 ************************************************************

NM_152232.1      ISWHTVNNTIPMSMCSKRCQSGQKKKPVGIHVCCFECIDCLPGTFLNHTEDEYECQACPN      540
NM_152232.4      ISWHTINNTIPMSMCSKRCQSGQKKKPVGIHVCCFECIDCLPGTFLNHTEDEYECQACPN      540
NM_152232.5      ISWHTINNTIPMSMCSKRCQSGQKKKPVGIHVCCFECIDCLPGTFLNHTEDEYECQACPN      540
NM_152232.6      ISWHTINNTIPMSMCSKRCQSGQKKKPVGIHVCCFECIDCLPGTFLNHTEDEYECQACPN      540
                 *****:******************************************************

NM_152232.1      NEWSYQSETSCFKRQLVFLEWHEAPTIAVALLAALGFLSTLAILVIFWRHFQTPIVRSAG      600
NM_152232.4      NEWSYQSETSCFKRQLVFLEWHEAPTIAVALLAALGFLSTLAILVIFWRHFQTPIVRSAG      600
NM_152232.5      NEWSYQSETSCFKRQLVFLEWHEAPTIAVALLAALGFLSTLAILVIFWRHFQTPIVRSAG      600
NM_152232.6      NEWSYQSETSCFKRQLVFLEWHEAPTIAVALLAALGFLSTLAILVIFWRHFQTPIVRSAG      600
                 ************************************************************

NM_152232.1      GPMCFLMLTLLLVAYMVVPVYVGPPKVSTCLCRQALFPLCFTICISCIAVRSFQIVCAFK      660
NM_152232.4      GPMCFLMLTLLLVAYMVVPVYVGPPKVSTCLCRQALFPLCFTICISCIAVRSFQIVCAFK      660
NM_152232.5      GPMCFLMLTLLLVAYMVVPVYVGPPKVSTCLCRQALFPLCFTICISCIAVRSFQIVCAFK      660
NM_152232.6      GPMCFLMLTLLLVAYMVVPVYVGPPKVSTCLCRQALFPLCFTICISCIAVRSFQIVCAFK      660
                 ************************************************************

NM_152232.1      MASRFPRAYSYWVRYQGPYVSMAFITVLKMVIVVIGMLATGLSPTTRTDPDDPKITIVSC      720
NM_152232.4      MASRFPRAYSYWVRYQGPYVSMAFITVLKMVIVVIGMLATGLSPTTRTDPDDPKITIVSC      720
NM_152232.5      MASRFPRAYSYWVRYQGPYVSMAFITVLKMVIVVIGMLATGLSPTTRTDPDDPKITIVSC      720
NM_152232.6      MASRFPRAYSYWVRYQGPYVSMAFITVLKMVIVVIGMLATGLSPTTRTDPDDPKITIVSC      720
```

```
                    ************************************************************

NM_152232.1         NPNYRNSLLFNTSLDLLLSVVGFSFAYMGKELPTNYNEAKFITLSMTFYFTSSVSLCTFM         780
NM_152232.4         NPNYRNSLLFNTSLDLLLSVVGFSFAYMGKELPTNYNEAKFITLSMTFYFTSSVSLCTFM         780
NM_152232.5         NPNYRNSLLFNTSLDLLLSVVGFSFAYMGKELPTNYNEAKFITLSMTFYFTSSVSLCTFM         780
NM_152232.6         NPNYRNSLLFNTSLDLLLSVVGFSFAYMGKELPTNYNEAKFITLSMTFYFTSSVSLCTFM         780
                    ************************************************************

NM_152232.1         SAYSGVLVTIVDLLVTVLNLLAISLGYFGPKCYMILFYPERNTPAYFNSMIQGYTMRRD 839
NM_152232.4         SAYSGVLVTIVDLLVTVLNLLAISLGYFGPKCYMILFYPERNTPAYFNSMIQGYTMRRD 839
NM_152232.5         SAYSGVLVTIVDLLVTVLNLLAISLGYFGPKCYMILFYPERNTPAYFNSMIQGYTMRRD 839
NM_152232.6         SAYSGVLVTIVDLLVTVLNLLAISLGYFGPKCYMILFYPERNTPAYFNSMIQGYTMRRD 839
                    ***********************************************************
```

**Figure S2:** Sequence alignment of human T1R2 RefSeq versions NM_152232.1, NM_152232.4, NM_152232.5 and NM_152232.6.
